# The Human Infertility Single-cell Testis Atlas (HISTA): An interactive molecular scRNA-Seq reference of the human testis

**DOI:** 10.1101/2023.09.23.558896

**Authors:** Eisa Mahyari, Katinka A. Vigh-Conrad, Clément Daube, Ana C. Lima, Jingtao Guo, Douglas T. Carrell, James M. Hotaling, Kenneth I. Aston, Donald F. Conrad

## Abstract

**Background:** The Human Infertility Single-cell Testis Atlas (HISTA) is an interactive web tool and a reference for navigating the transcriptome of the human testis. It was developed using joint analyses of scRNA-Seq datasets derived from a dozen donors, including healthy adult controls, juveniles, and several infertility cases. HISTA is very different than other websites of testis scRNA-seq data, providing visualization and hypothesis testing tools on a batch-removed and integrated dataset of 23429 genes measured across 26093 cells using.

**Objective:** The main goal of this manuscript is to describe HISTA in detail and highlight its unique and novel features.

**Methods:** Therefore, we used HISTA as a guide for its application and demonstrated HISTA’s translational capacity to follow up on two observations of biological relevance.

**Results:** Our first analytical vignette identifies novel groupings of tightly regulated long non-coding RNA (lncRNA) molecules throughout spermatogenesis, suggesting specific functional genomics of these groupings. This analysis also found highly controlled expression of pairs of sense and antisense transcripts, suggesting conjoined regulatory mechanisms. In the next investigative vignette, we examined gene patterns in undifferentiated spermatogonia (USgs). We found the NANOS family of genes function as key drivers of transcriptomic signatures involved in human spermatogonial self-renewal programming; for the first time, demonstrating the relationship of NANOS1/2/3 transcripts in humans with scRNA-seq.

**Discussion and Conclusions:** Using HISTA, we found new observations that contribute to unraveling the mechanisms behind transcriptional regulation and maintenance germ cells across spermatogenesis. Furthermore, our findings provide guidance on future validation studies and experimental direction. Overall, HISTA continues to be utilized in testis-related research, and thus is updated regularly with new analytical methods, visualizations, and data. We aim to have it serve as a research environment for a broad range of investigators looking to explore the testis tissue and male infertility.

**Availability and Implementation:** HISTA is available as an interactive web tool: https://conradlab.shinyapps.io/HISTA

Source code and documentation for HISTA are provided on GitHub: https://github.com/eisascience/HISTA

## INTRODUCTION

The testis is a complex reproductive tissue with unique microenvironments and cell types that recently has been the focus of numerous single-cell transcriptomics (scRNA-Seq) studies to unravel both normal and pathological features ^1–7^. Recently, we published our findings ^1^ with 26093 cells, derived from testis biopsies of juveniles, normal adults, and adults with known infertility (azoospermia, ejaculatory dysfunction, and Klinefelter Syndrome). To facilitate our research of the testis transcriptomic data at the core of our work, we developed the Human Infertility Single-cell Testis Atlas (HISTA). HISTA is an interactive reference and a rich resource for testicular biology (Fig 1), already utilized in publications ^1,8^ and internal reports supporting translational reproductive biology. There are two general ways to use HISTA: A) as a web tool to explore testis scRNA-Seq data (https://conradlab.shinyapps.io/HISTA/) or B) as a resource to download for analyses that generate or test hypotheses. In this manuscript, in our analytical vignettes, we demonstrate both applications.

**Fig 1:**
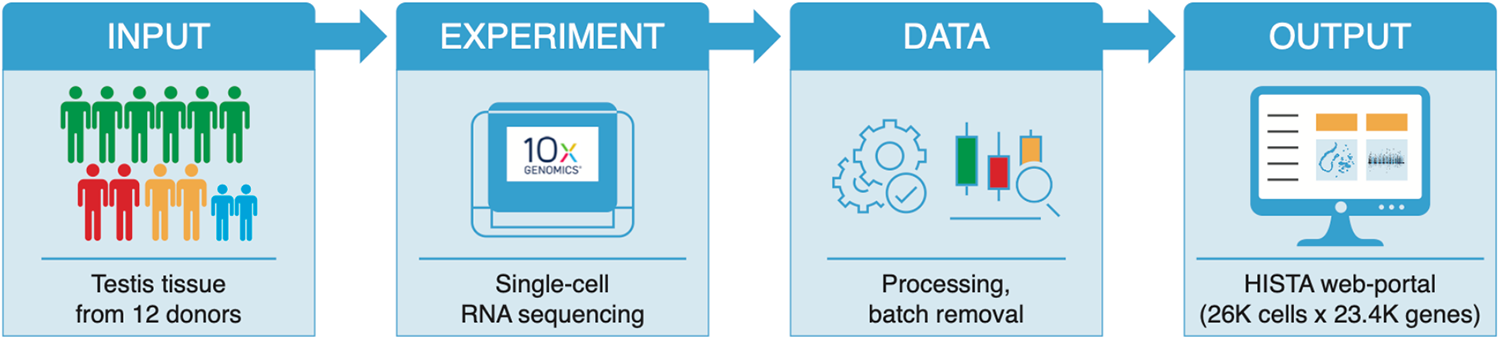
The Human Infertility Single-cell Testis Atlas (HISTA) web-portal (2022 release) (https://conradlab.shinyapps.io/HISTA/) enables exploration and hypothesis testing across 26093 high-quality cells derived from testis biopsies (2 juveniles, 6 normal adults, 1 adult with azoospermia, 1 adult with ejaculatory dysfunction, and 2 adults with Klinefelter Syndrome (KS)).

Other websites that allow interaction with scRNA-seq data of human testis samples have been recently summarized ^9^. None provided a multi-dataset integrated resource carefully curated to minimize technical noise and batch effects and none identified functional clusters of genes across the various testis cell types and spectra of states. Furthermore, none provide an interpretable, scalable, and generalizable approach to interrogate and characterize new omics data. Several of the reported websites ^9^, are simply post-processed data on the UCSC cell browser. The Human cell landscape (https://bis.zju.edu.cn/HCL) houses the data from a recent pan tissue study ^10^ which unfortunately excluded the testis, so this atlas utilizes another previously published dataset ^6^, as their testis dataset (https://bis.zju.edu.cn/HCL/dpline.html?tissue=Testis). Websites like PanglaoDB ^11^ and ReproGenomics ^12^ are great resources for exploring and downloading individual datasets; however, cross-dataset integration is a current feature. The Germline Atlas (https://germline.mcdb.ucla.edu/) is another resource that houses *in vitro* and *in vivo* data on specific developmental stages. More recently, a website by our collaborators (https://humantestisatlas.shinyapps.io/humantestisatlas1/), enables general exploration of individually processed data of their “puberty atlas”, “young adult atlas”, or “adult SSC states”. Lastly, Deeply Integrated human Single-Cell Omics (DISCO) (https://www.immunesinglecell.org/) is a highly efficient and generalizable framework with a comprehensive set of human tissues with the capacity to integrate a limited number of datasets for downstream analysis. DISCO currently features about 40 QC passing samples (of 60 total) for the testis. Their current atlas (V1.0) has focused mainly on early human life from early weeks of gestation to infancy and adulthood (GSE161617, GSE143381, GSE124263, GSE120508, GSE149512 and E-MTAB-10551); GSE120508 has 3 adults and 2 juvenile that overlap with HISTA. DISCO provides interaction to search gene expression on a 2D projection across several metadata options. Although their data is integrated, it still appears batch is a driver of some of the clusters; HISTA was manually and carefully curated to minimize such unwanted noise. More importantly, DISCO lacks a full spectrum of germ cells representing the entire spermatogenesis trajectory. Therefore, HISTA remains the only interactive tool with a comprehensive representation of spermatogenesis, focused on the pathology of infertility as well as development, curated with multiple types of analyses and figures to provide translational tools examining the transcriptomic landscape of the testis. Furthermore, using HISTA’s data, specifically the SDA ^13^ model, new data (single-cell or spatial omics) can be annotated by transferring the components fast and relatively simple; the supplemental section on SDA describes specifically the application of SDA to scRNA-seq data ^1,2^ and the transfer of HISTA’s model to new data.

In this manuscript, our main objective is to broaden and enhance the usage of HISTA by discussing the technical details of its construction and application through two translational vignettes pertaining to regulating and maintaining human germ cells. Therefore, we organize the remainder of this manuscript to have separated sections for each analysis. In the first vignette, we investigate and report on novel sets of long non-coding RNA (lncRNA) molecules that tightly regulate various transcriptional stages of spermatogenesis. We then focus on Undifferentiated Spermatogonia (USgs), leveraging our unique amalgam dataset to find new functional characteristics and transcriptional signatures. We additionally hope that these computational investigations highlight how HISTA is a valuable resource for the research community and how to utilize HISTA best to enable hypothesis generation, as well as follow-up and validation studies to understand testicular biology and infertility.

### Vignette 1: Novel sets of lncRNA molecules identified as key regulatory processes in spermatogenesis with potential implications for fertility

There are at least 58,000 Long non-coding RNAs (lncRNAs) in the human genome that are known to have specific expression patterns by cell type and tissue, yet their full spectrum of function remains elusive ^14,15^. The pathologies that mechanistically have been linked to lncRNAs include cardiovascular disease, diabetes, Alzheimer’s, Parkinson’s, and autoimmune disease. The testis, an amalgam of somatic cells with continuously developing germ cells, provides a unique opportunity to uncover novel roles and mechanisms of lncRNA.

Several studies have reported on the functional mechanisms of lncRNA ^16–18^ in various tissues. Several others have specifically focused on the testis ^19–22^, including studies in domesticated animals linking fertility robustness to several lncRNAs ^23–25^. In a study with mice using microarrays, the testis was shown to have the highest number of expressed and specific lncRNAs ^22^. More recently, using public microarray data, Joshi et al. identified fifteen functionally conserved lncRNAs across multiple studies in humans and mice ^25^, which with HISTA, we explore deeply. Broadly, in the testis, lncRNA could A) regulate the expression of protein-coding genes involved in sperm production or function ^26^. B) Directly interact with sperm cells or other cells in the testis ^27^, potentially binding to receptors on the surface of sperm cells, altering their motility or ability to fertilize an egg or interact with the Sertoli cells or Leydig cells. C) Influence the hormonal environment of the testis ^28,29^; binding to and regulating the activity of hormones, such as testosterone or follicle-stimulating hormone. D) Change the development or maintenance of the blood-testis barrier ^30^. E) Affect the microenvironment of the testis by influencing the production or function of extracellular matrix molecules, such as collagen or proteoglycans, that provide structural support for sperm cells ^27,31^. F) Induce changes in immune responses in the testis ^27,32^, either to regulate the production of inflammatory molecules or to regulate the activity of immune cells involved in the clearance of abnormal or damaged sperm cells. Herein, we present novel groupings of lncRNAs that, individually, the majority have been previously reported as testis-specific. The main novelty herein is the grouping of these sets, which represent coexpressed or coregulated clusters of these lncRNAs, grouped with other transcripts found in specific cells in various transcriptional stages of germ cells from spermatogonia to spermatids.

### Vignette 2: A Cyclical transcriptional profile describing maintenance and regulation of early spermatogonial populations

Undifferentiated spermatogonia (USgs) are a critical population of germ cells with stem properties that maintain their populations and feed spermatogenesis. However, across various single-cell studies of the testis, there are inconsistencies in identifying and labeling these cells. For example, some studies focus on a high-scope population label such as spermatogonia ^33,34^, whereas others have more refined sub-populations and unique naming structures ^5,35,36^. To add to the complexity, histologically, SSC can be split in humans into Apale (Ap) and Adark (Ad) spermatogonia ^37^, which are not distinct populations transcriptionally. To unify and shed clarity, we review key findings from scRNA-seq papers that discuss USgs.

● Hermann et al.’s analysis of human cells shows three major populations across Spermatogonial cells: the SSCs, progenitors, and Differentiating. Using pseudotime analysis, they show *ID4* and *GFRA1* expressed in early SSCs. *NANOS3* is found mid-way, and *STRA8* is found towards the differentiating Sgs ^7^.
● Guo et al., by analyzing just spermatogonial cells, describe five discrete transcriptional states for SSCs; State 0 through 4. They Identify *PIWIL4, PPP1R36, EGR4, MSL3,* and *TSPAN33* as genes specifically expressed in State 0. They Identify *ID4, UTF1, L1TD1,* and *ETV5*, to be enriched in States 0 and 1, *MKI67* in States 2 and 3, *STRA8* in State 4, and *MAGEA4* and *DMRT1* as expressed across all States ^6^.
● Guo et al., in a later publication, identified a trajectory of germ cells labeled as USgs, Differentiating Sgs, spermatocytes, and spermatids. Their findings show *UTF1, PIWIL4, TSPAN33,* and *GFRA1* to be expressed specifically in the USgs cluster, and their results show this population is the most frequent germ cell population prior to puberty ^5^.
● In a follow-up study, using embryonic, fetal, and postnatal samples, the same research group proposes a transcriptional trajectory for the development of human primordial germ cell (PGC) to adult Sgs, respectively as Embryo-Fetal, Fetal-Infant, State0, State1, State2, State3, and State4. They identify *NANOG*, *POU5F1*, and *DPPA3* as highly specific to the Embryo-Fetal cluster. *PIWIL4*, *EGR4*, and *MSL3* are specific to the Fetal-infant & State0 clusters. *GFRA1* is reported to be specific to State1. The other states, State2, 3, & 4 are not shown to have specific expression, although they exhibit proliferation signals such as *MKI67* expression ^4^.
● Shami et al. investigated 3 mammalian species (human, *Rhesus* macaque, mouse) testis and defined 6 spermatogonial populations labeled SPG1 through SPG6. Humans (and macaque) show *MORC1, MSL4, TCF3, FMR1, MAGEB2,* and *CDK17* as SPG1-specific transcripts. SPG2-3 express *L1TD1*, followed by *KIT* and *DMRTB1* in SPG4-6, and *STRA8* mainly in SPG5-6.
● Lau et al. studied the *Cynomolgus* macaque testis. Their high-level pan testis spermatogenesis trajectory resembles the human data in HISTA, including the ring structure defined by spermatogonial cells. When they reprocessed their spermatogonial cells, this cyclical pattern persisted; however, they suggest that the order in these populations seen on their 2D projection is not the biological order. Their groupings include Undiff1/2/3, E-diff1/2/3/4, Mid-diff, late-diff, pre-Lep, and Lep, respectively. They support their clusters by showing differentially expressed genes. Of these genes, *DNMT3L* and *EGR4* are enriched most in Undiff1 and are absent in the later populations. Conversely, *STRA8* and *REC8* are enriched in pre-Lep and Lep but lacking in the early populations. As in humans, *NANOS3* was absent in early Undiff1/2/3 and later pre-Lep and Lep populations but enriched mainly in E-diff3-4.

A common finding across these studies highlights transcriptional patterns such as *ID4* and *GFRA1* being expressed in early undifferentiated transcriptional stages of spermatogonial development, *NANOS3* in mid-way stages, and *STRA8* in differentiating spermatogonia. Given that the integrated data of HISTA is an amalgam of several of the studies described above, it innately has higher precision and statistical power to evaluate the expression of the genes described, which can be found in the results part of our second vignette. A repeating pattern from the above findings that complement our findings is that the NANOS family of genes are differentially expressed in USgs populations and are critical to a cyclical signal of USgs renewal.

Humans have three NANOS genes orthologous to the gene *nanos* in drosophila melanogaster. In flies, *nanos* is required for male fertility by preventing all germline stem cells from differentiation ^38^. *NANOS1* is thought to inhibit germ cell apoptosis in humans ^39^, but how crucial it is for USgs, is unknown, as knockout mice for Nanos1 are not only viable, they are fertile ^40^. *NANOS2*, on the other hand, appears to function in the sexual differentiation of germ cells during early embryonic stages and after birth; in mice, the transcriptional expression is lowered after the E16.5 ^41^. Therefore, *NANOS2* is thought to function in maintaining male embryonic gonocytes ^42^ and lowering the proliferation rate of USgs by targeted degradation of specific RNA molecules ^43^. Functionally, *NANOS3* is thought to inhibit spermatogonial differentiation and thus maintain the undifferentiated population ^44^. At the single-cell resolution, only *NANOS3* has been previously described ^45^ as a primordial germ cell (PGC) expressed gene in humans, and more recently, it is expressed in early germ cells of the *Cynomolgus* macaque testis ^35^. NANOS1 and NANOS2 proteins have also been previously characterized as conserved in the testis ^46^. Lastly, this family of genes may have additional roles in other tissues and other cell types of the testis tissue; for example, *NANOS3* in sheep has also been identified as a regulator of Leydig cell (LC) testosterone production ^47^.

## MATERIAL AND METHODS

### Data acquisition and preprocessing

As reported previously, HISTA contains data from several published studies six adults (GSE109037 & GSE120508), two juveniles (GSE120506), an idiopathic azoospermic patient (INF1), two Klinefelter patients (KS1 and KS2), and a patient with secondary infertility treated as a control (INF2/CNT-U4) (all infertility cases in GSE169062). The raw count matrices of these datasets were combined and processed using several available tools as previously described ^1^.

### SDA and component annotation

The SDA run housed by HISTA was set to identify 150 components individually examined to identify cell type of action and transcriptional function ^1^, displayed in the information box across HISTA, including the ‘Main’ tab of HISTA. Beyond our previous description of how SDA was implemented for HISTA ^1^, we elaborate on this machine-learning approach in detail in the supplemental section.

### Shiny App

HISTA is developed using the “shinydashboard” package ^48^ on the Shiny framework ^49^ written in R ^50^ with the help of several key packages, including ggplot^51^, dplyr ^52,53^, and data.table ^54^ as well as Seurat ^52^, which was used in parallel processing of this data as described previously^1^. The code is available on Github (https://github.com/eisascience/HISTA)^55^.

### lncRNAs

To identify if a given component is enriched or depleted of the lncRNAs (or any gene set from the user), a background distribution is obtained by repeated (N=3) random sampling of an equal number of (or at least 1000) genes. Because the randomly obtained sets depict a normal distribution, we can conservatively use the three standard deviations from the mean to identify components highly enriched for lncRNAs (Fig 2A), such that dashed lines identify the range representing 99.7% of the data. Next, to compare the similarity of germ cell-associated components with regard to lncRNAs found in each, we computed Pearson’s correlation coefficients from an identity matrix of these genes across the SDA loadings with hierarchical pair-wise clustering (Fig 2B).

**Fig 2.**
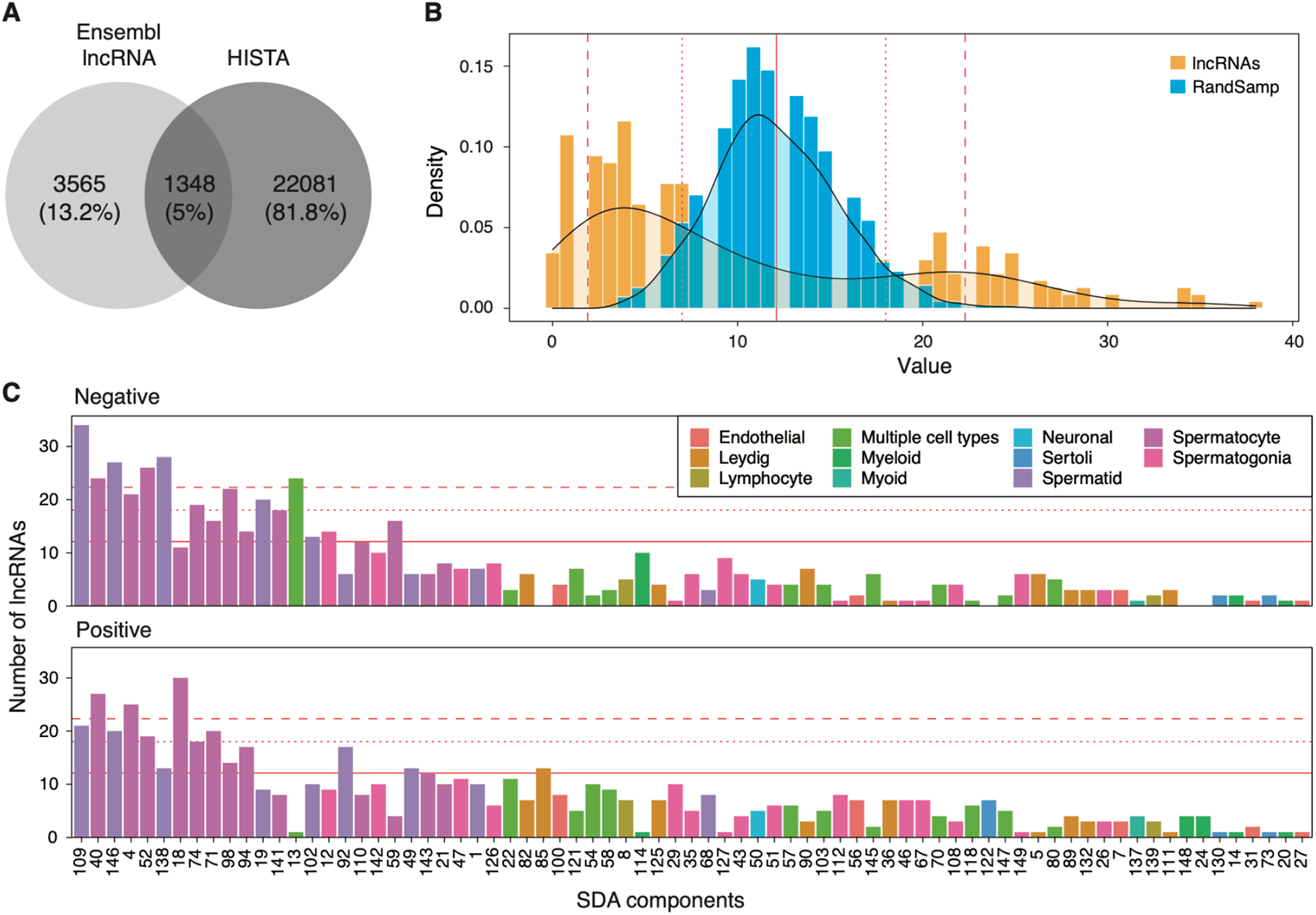
**A**) Overlap of lncRNA genes (annotated on the Ensembl reference genome using the biomaRt package ^68^) with those found in HISTA. **B)** The overlap distribution of lncRNA genes versus simulated repeated (N=3) random sampling of an equal number of genes (N=1348) overlapping with the top 200 positive and negative loaded genes across SDA components. The solid line identifies the mean, the dashed lines represent the mean +/-3sd boundary and the dotted lines represent the 5 to 95 percentile range of the random sample distribution, to guide finding components enriched or depleted of lncRNAs. **C)** Enrichment of SDA components that passed QC (i.e., not removed as batch or noise) by lncRNA genes, colored by the major cell type each component highly scores. The solid line represents the average (12.1) number of genes overlapping each SDA component across all components in the simulated random sample set of genes (see Fig 1 B). The dotted line (18) and the dashed line (22.3) are the 95% quantile and mean + 3 x standard deviation of this random set simulated overlap to guide as a positive threshold for enrichment.

In the supplement, we also report additional findings pertaining to sense and antisense expression patterns, which are projected across pseudotime (spermatogenesis trajectory) for each pair. A simple subtraction of the antisense from the sense expression is also shown as a simplistic measure of a delta expression.

### USgs

HISTA browser and its data were used for the analyses in this manuscript and the production of the figures, including those pertaining to USgs. As with most of this manuscript, we utilized HISTA for analysis and visualization, although many have been externally faceted and organized with image processing software for publication. However, as a parallel analytical approach to SDA and how HISTA represents USgs, using the HISTA data, we performed a computational validation of our results, reported as supplemental results. In R, we subset to the Usgs cells of control adults (representing the donut and its neck structures) and reprocessed using Seurat ^52^ to obtain the top 500 variable genes. This set was used to compute K-means clustering with K heuristically set to provide equal-size clusters. The clusters were then used to train a random-forest classifier (80% training 20% testing scheme) scheme, yielding an importance ranking of genes using the Gini index. Broadly, these steps intend to provide a parallel approach to SDA in identifying clusters of cells in an unbiased fashion and the most important genes driving those clusters. Since USgs, like all other cells, are indeed a spectrum, we used Scorpius ^56^ to determine the gene expression pattern of the pseudotime through the K-means clusters identified (Sup Fig 2).

## RESULTS

Navigation of HISTA is through a tabular set of analyses, starting with the ‘Main’ tab as the entry point. However, for those with an R (or other coding) environment, the data object can be downloaded for deeper analysis and customization of visualizations. In supplement to this manuscript, we provide a detailed guide on each tab. However, to highlight specific features, translational capacity, and possible navigation of HISTA, we provide two analytical vignettes broadly related to transcriptional regulation and maintenance of germ cells, which would interest the research community.

### Vignettes 1

In our first vignette, we follow up on an observation that several of HISTA’s components were enriched with long non-coding RNAs (lncRNAs). Navigating to HISTA’s lncRNA tab, we find that 38% of annotated lncRNAs (hg19 genome assembly) show evidence of expression in the human testis transcriptome (Fig 2A). These lncRNAs show far greater co-regulation than expected due to chance. Thus, they are more strongly enriched in germ cell SDA components than somatic cell components (Figs 2-4, Tables 1-2). On average, there are ten lncRNAs per SDA component, with a standard deviation (sd) of 8.9, and 12 random genes, with much smaller variance (sd = 3.4), which enabled defining thresholds to identify components enriched for or depleted of lncRNAs (Table 1). Since the cell type annotation of each component has been established as an estimate of the major cell type(s) or specific functions of those cells, we demonstrate that the components most enriched with lncRNAs are components pertaining to specific germ cells. Specifically, spermatocytes and Spermatids have the most lncRNA transcription (left to right in Fig 2C).

**Table 1:** Summary statistics, evaluating enrichment of lncRNA genes by enumerating gene overlap across the components in contrast to repeated (N=3) random sampling of an equal number of genes (N=1348). Mean = average number of lncRNAs or random protein coding with high loading on an SDA component. The average here is the average over all QC-passed SDA components.

| Geneset | Mean ($\mu$) | Sd ($\sigma$) | $\mu + 3\sigma$ | $\mu - 3\sigma$ | 5% | 50% | 95% | CV |
| --- | --- | --- | --- | --- | --- | --- | --- | --- |
| lncRNA | 10.5 | 8.9 | 37.2 | -16.2 | 1 | 7 | 27 | 0.84 |
| Random Sample | 12.1 | 3.4 | 22.3 | 1.9 | 7 | 12 | 18 | 0.28 |

**Table 2.**
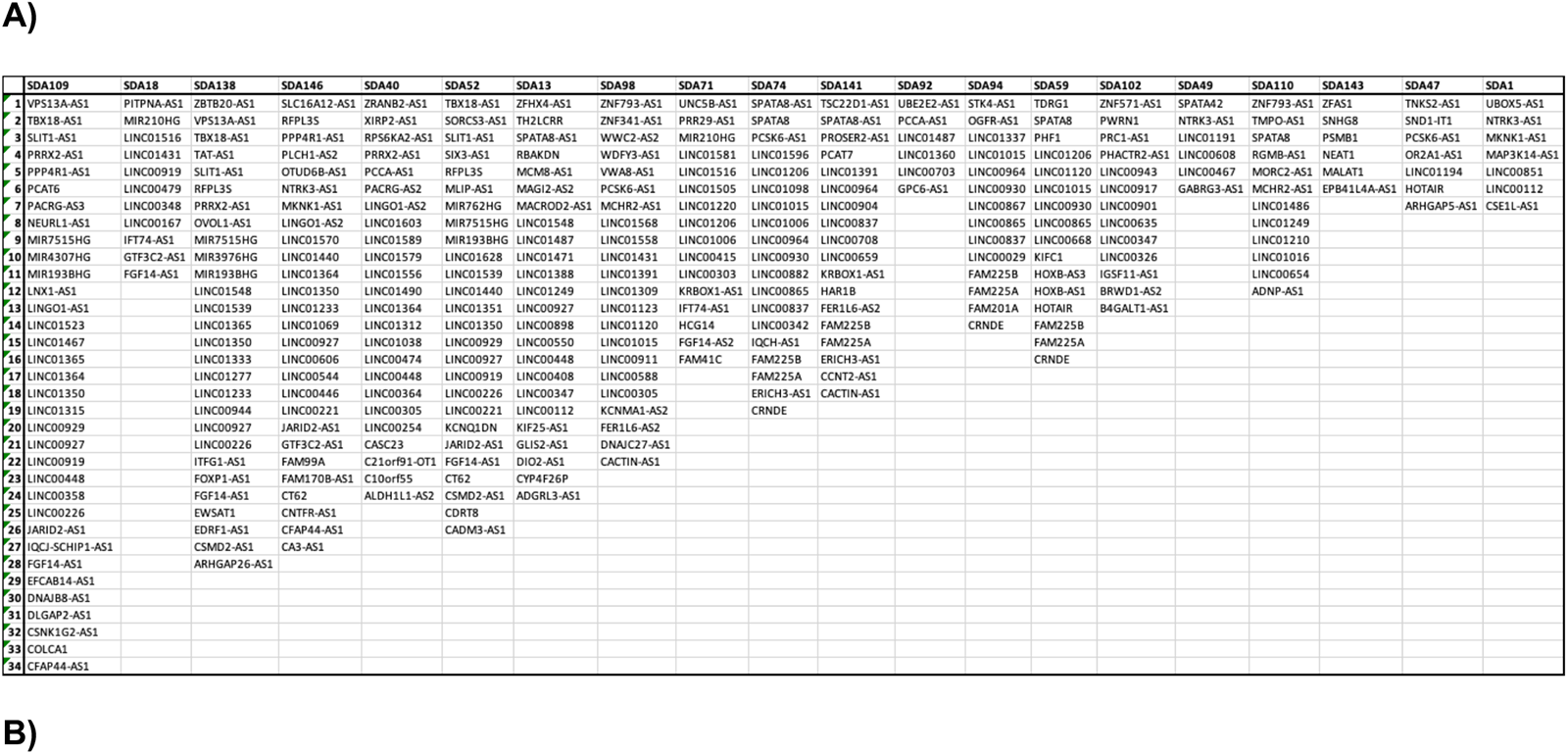
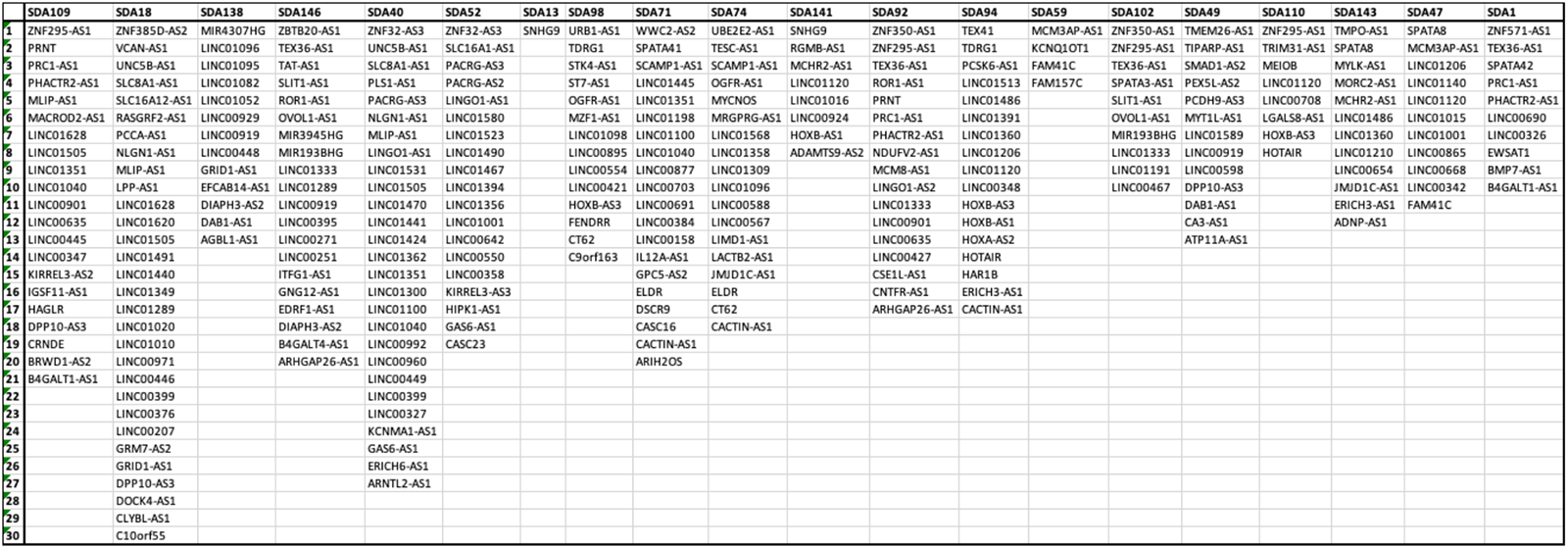
Top loaded lncRNA transcripts per component found across spermatogenesis in germ cells. **A**) Overlapping of available lncRNAs with top 200 positively loaded genes. **B**) Overlapping of available lncRNAs with top 200 negatively loaded genes. In both comparisons, the top 20 components that are enriched for lncRNAs and highly scored germ cells.

Across spermatogenesis, the identified lncRNA signatures identify specific populations of cells and differentially expressed groups of functionally related transcripts, including known genes, identifying biologically relevant signatures. For a global view, we compiled the complete set of the components scoring germ cells by directly accessing the data of HISTA (Fig 3); however, on the ‘Main’ tab of HISTA, the scoring pattern of any individual component colorized on a 2D representation can be found. As a case example, tabulated on ‘the lncRNA’ tab, component 109 (SDA109) has the most significant number of distinct lncRNA transcripts.

**Fig 3.**
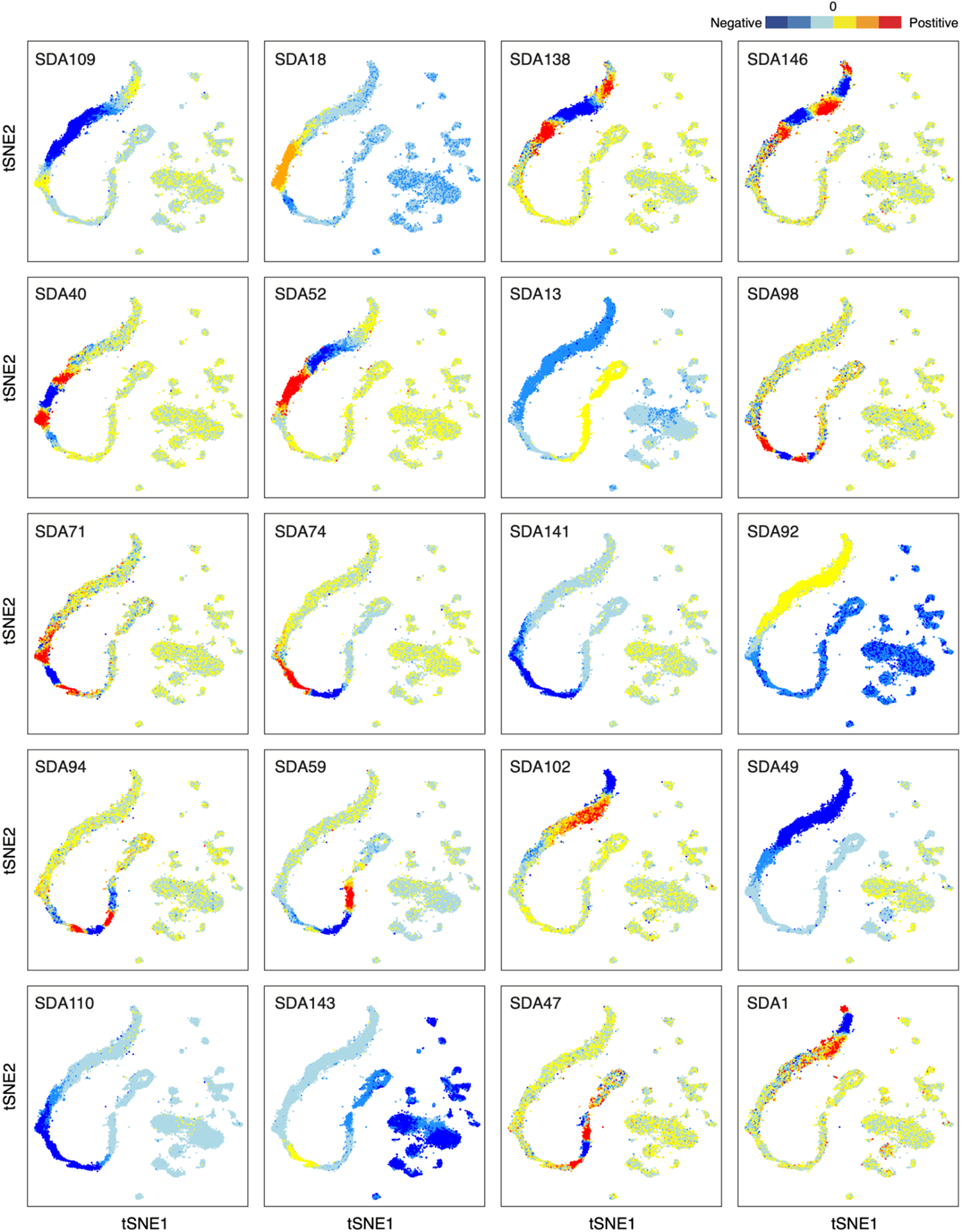
Cell scoring patterns of SDA components that are found highly enriched in lncRNA; grouped by the cell population they are most enriched in. Each component identifies differentiating functional sets of lncRNA expression signatures related to various cells and mechanisms of spermatogenesis. Using HISTA, each component can be investigated to identify corresponding cell types and pathologies for translational needs; see HISTA’s ‘Index of components’ tab, where these components have been described in detail. Broadly, the scores are a distribution that identifies low, mid, and high groupings. When curated as in this figure, an emergent feature is that the components are finding various scoring patterns, beyond what is quantitatively achievable by manual phenotyping or clustering with a generalized pipeline. Some components show rigid whereas others are more distributed in terms of which cells or populations they highlight e.g. SDA138 vs SDA109 respectively.

SDA109 identifies a unique signature of a population of cells in spermiogenesis that are scored negatively but flanked on both sides by strongly positive scoring cells. Such a differential pattern is essential when considering the expression patterns and, thus, the functional effect of the top-loading genes of this (or any) component. Searching the genes found in this component on the ‘Main’ tab, a logical pattern emerges that positively loaded genes are detected in the positive scoring cells but, lacking in the negative scoring cells and vice-versa for negative loaded genes; this is because SDA109 score, like many other component scores, is distributed around zero.

Using the HISTA’s ‘lncRNA’ tab, we can identify the top-loaded lncRNAs of SDA109 include *DLGAP2-AS1, PPP4R1-AS1, LINC01523, MIR4307HG, LINC00448, LINC00358, LINC00226, JARID2-AS1, LNX1-AS1, LINC00929, LINC01365, FGF14-AS1, VPS13A-AS1, PRRX2-AS1, MIR7515HG, TBX18-AS1, MIR193BHG, SLIT1-AS1, LINC00927, LINC00919, PACRG-AS3, LINGO1-AS1, LINC01315, CFAP44-AS1, LINC01364, LINC01467, NEURL1-AS1, DNAJB8-AS1, CSNK1G2-AS1, LINC01350, COLCA1, IQCJ-SCHIP1-AS1, EFCAB14-AS1,* and *PCAT6*. HISTA’s built-in gene ontology (GO), on the ‘Main’ tab, suggests SDA109 has a functional association with spermiogenesis and acrosome development and regulation. Furthermore, many of the top-loaded genes of this component map to spermatid flagellum development and motility (*TRIM42, TTLL2, DCDC2C, CCDC179, PRSS58, C10orf120, FAM71B, C20orf173, TEKT5, DNAJB7, CCDC185, TP53TG5, LRRC72, FAM24A, TULP2, LRRC3B*) as well as other testis-specific genes, but with unknown or non-germ cells specific functions (*RAB21, MARCH8, IQCF6, MSMP, ELL, C17orf50, NRDC, GGTLC1*). It should be noted that these genes are found together, implying some correlation relative to the magnitude and direction of the gene loadings themselves specific to the cells that scored most in either direction of the score distribution. Simply put, SDA109 as an example, uniquely identifies a population of spermatids with a transcriptional signature driven by the top-loaded genes.

Another lncRNA-enriched component of interest is component 52 (SDA52), which functionally maps to sperm motility, flagella development, and sperm capacitation. SDA52 functionally is related (not identical) to SDA109 as there is an overlap in cells scored as non-zero and overlapping top-loaded genes. Specifically, we find that the cell score pattern of SDA52 identifies a boundary between spermatocytes in meiosis II and the round spermatids (Fig 3). Furthermore, a unique set of top-loaded genes, including lncRNAs, were found in SDA52 and not SDA109 (Table 2). These results suggest that although global transcriptional patterns govern spermatogenesis locally, at each transcriptional stage, there are specialized mechanisms of control involving specific lncRNAs.

To quantify and determine the specificity of lncRNAs captured by the SDA components globally, we searched the top-loaded lncRNAs across them and found 365 unique lncRNAs (Table 2). Using the HISTA dataset, these lncRNAs’ expressions were found to be differentially expressed at specific transcriptional stages of spermatogenesis (Fig 4A). Each transcriptional stage of spermatogenesis has a unique signature, but the most extensive set of uniquely expressed lncRNAs is found in the pachytene-diplotene spermatocytes, followed by round spermatids. Since there is some functional overlap across these lncRNA enriched components, we performed correlation analysis with the SDA loadings (Fig 4B); a similar figure for the 365 lncRNAs or any other gene sets can be computed within HISTA using the ‘Top loaded components’ tab to identify functionally similar components. Although the correlation coefficients we identified are moderate to low, components with similar lncRNA loading tend to score the same cell populations suggesting lncRNA and population specificity. For example, components SDA143 and SDA110 which cluster together, are expressed in meiotically dividing spermatocytes, or components SDA109, SDA52, and SDA138, which score the round spermatids (Fig 3).

**Fig 4.**
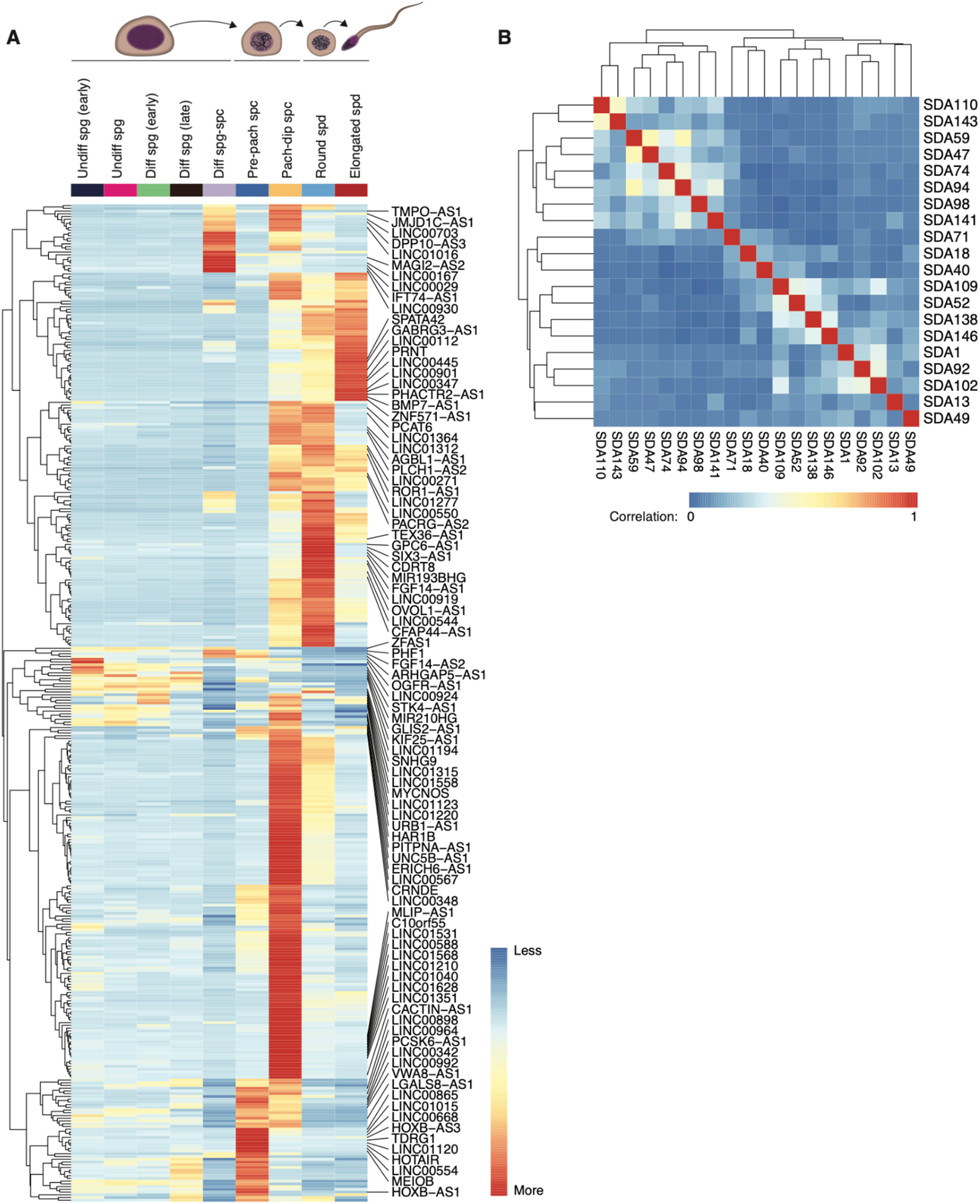
**A)** Relative average lncRNA expression of control donors only, columns ordered by spermatogenesis trajectory, identifies differential lncRNA usage across stages of spermatogenesis. Shown here are the union of all lncRNAs identified within the top 200 gene loadings of any SDA component. **B)** Identification of similarity amongst SDA components using correlation analysis on an “identity matrix”. By computing Pearson’s correlation on a boolean matrix of 0-values or 1-values, where the latter identifies a particular lncRNA is found in a given SDA component; the pair-wise hierarchical clustering groups the similar components. This analysis uses the same set of lncRNAs as in **A)**.

While investigating these components such as SDA109, we found several of them to be also highly enriched for antisense genes, often paired with the sense counterpart in the same component. We followup on this observation in the supplement section (Supp Fig 1). Briefly, we find these antisense transcripts to be highly regulated in expression across spermatogenesis, and their expression pattern relative to the matching sense transcripts plays a role in regulating the transcriptional mechanisms.

### Vignettes 2

Our second analysis utilized HISTA to evaluate a hypothesis about spermatogonial stem cell (SSC) self-renewal. USgs are the earliest precursors of spermatogenesis and, in our samples, comprise about 10% [5-25%] of the total germ cells in healthy human adults ^1^. We initially observed undifferentiated spermatogonia (USgs) in a ring structure when projected in a 2-dimensional representation. Interestingly, we also found a similar torus structure of USgs in other publications of humans ^36^ as well as other mammals^33,35^. We examined the cyclical nature of USgs with two approaches, the first using the HISTA browser and the second using the dataset of HISTA as a computational validation of our findings using alternate methods.

Navigating the components, we identified four SDA components (SDA112, SDA126, SDA149, and SDA127) mapping to the USgs ring (Fig 5A). Each component provides a unique signature, which, relative to pseudotime on the germ cells, is depicted as a wave within the ‘Pseudotime meta’ tab of HISTA (Fig 5B). These components identified *NANOS2* and *NANOS3* as highly important (Fig 6A), which were found expressed in specific regions of the USgs structure (Fig 6B), alongside several other genes (Fig 7) as possible drivers of the maintenance and regulation of these USgs; what is novel in HISTA is indeed the grouping of genes within these components that identifies functional relationships.

**Fig 5:**
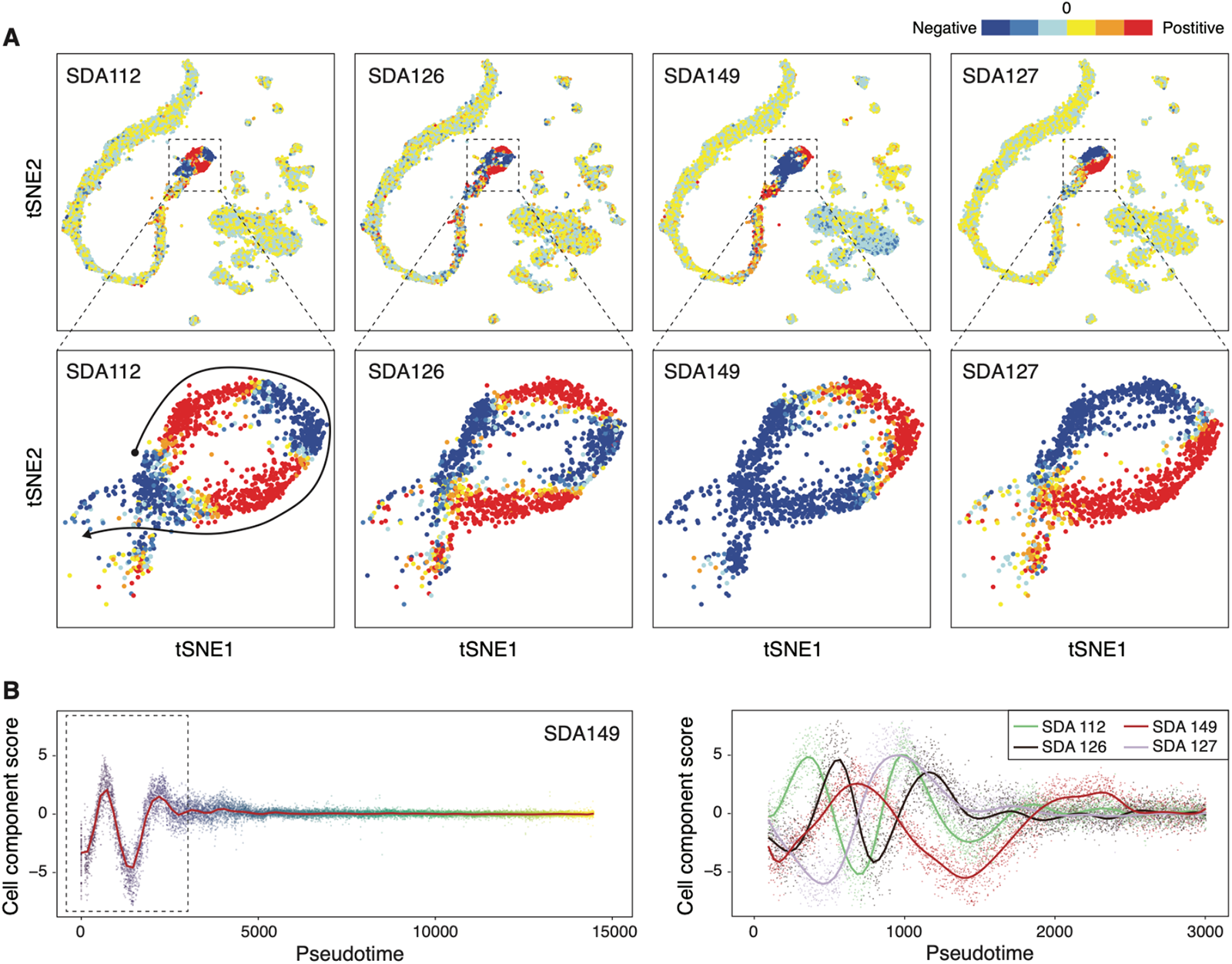
**A)** Four main components that score the undifferentiated spermatogonial cells differentially, showing structured gene expression patterns, and highlighting areas of high to low expression patterns. **B)** The patterns in which SDA components score cells, projected across pseudotime-ordered germ cells, has been previously described and shown informative in describing genomic mechanisms and expression patterns ^1,2^. Herein, we are focused on the SDA components that are highly variable only at the early stages of the spermatogenesis pseudo-trajectory; thus in the right figure, we zoom to this region on pseudotime. The observed pattern is that initially, as SDA112 rises, it is followed by SDA126 and SDA149, but SDA127 decreases. Later in the second wave, SDA127 reaches a maxima first, followed by SDA112 and SDA126, but at the same time, SDA149 decreases.

**Fig 6:**
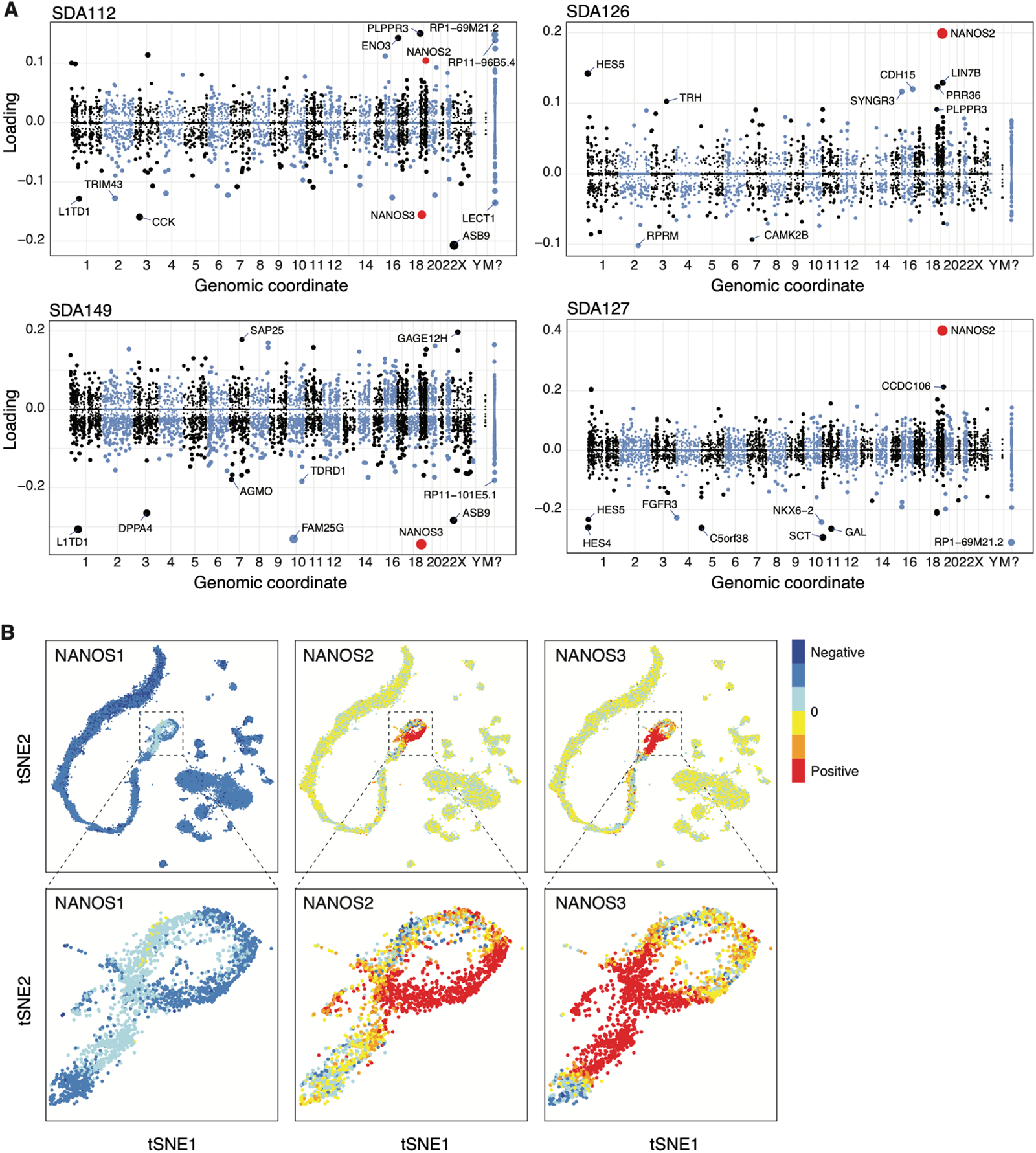
**A)** Top loaded genes of SDA components 112, 126, 149, and 127, obtained from HISTA, pertaining to USgs showing NANOS2/3 are highly loaded in these components. **B)** Expression of NANOS 1, 2, and 3 in early spermatogonia. SDA112 highlights transcriptomic mechanisms that target the USgs in a unique pattern. *NANOS2* and *NANOS3* are in the top-loaded genes and are in opposite loading directions, suggesting individual roles for each of these genes. The expression of *TRIM43, TRIM51, CCK, CCL17, SLC22A1*, and *CWH43*, found in the negatively loaded genes, have a very precise and localized expression within the USgs ring structure (Fig 7), suggesting a level of orchestration functionally. SDA126 is a rotational shift in which cells score positive relative to SDA112. *NANOS2* is SDA126’s top positively loaded gene, followed by *HES5*. SDA149 scores spermatogonia differentially. In the ‘pseudotime’ tab of HISTA, the germ cell scores ordered by pseudotime identify a wave pattern that decays as the germ cells enter meiosis. This component has *NANOS3* as the most negatively loaded gene in this component. SDA127 splits the spermatogonia in the ring structure into two sections, almost 90 degrees relative to the slices of SDA149. Interestingly, *NANOS2* is identified as the top positive loaded gene, which is very specifically expressed in the positively scored cells.

**Fig 7:**
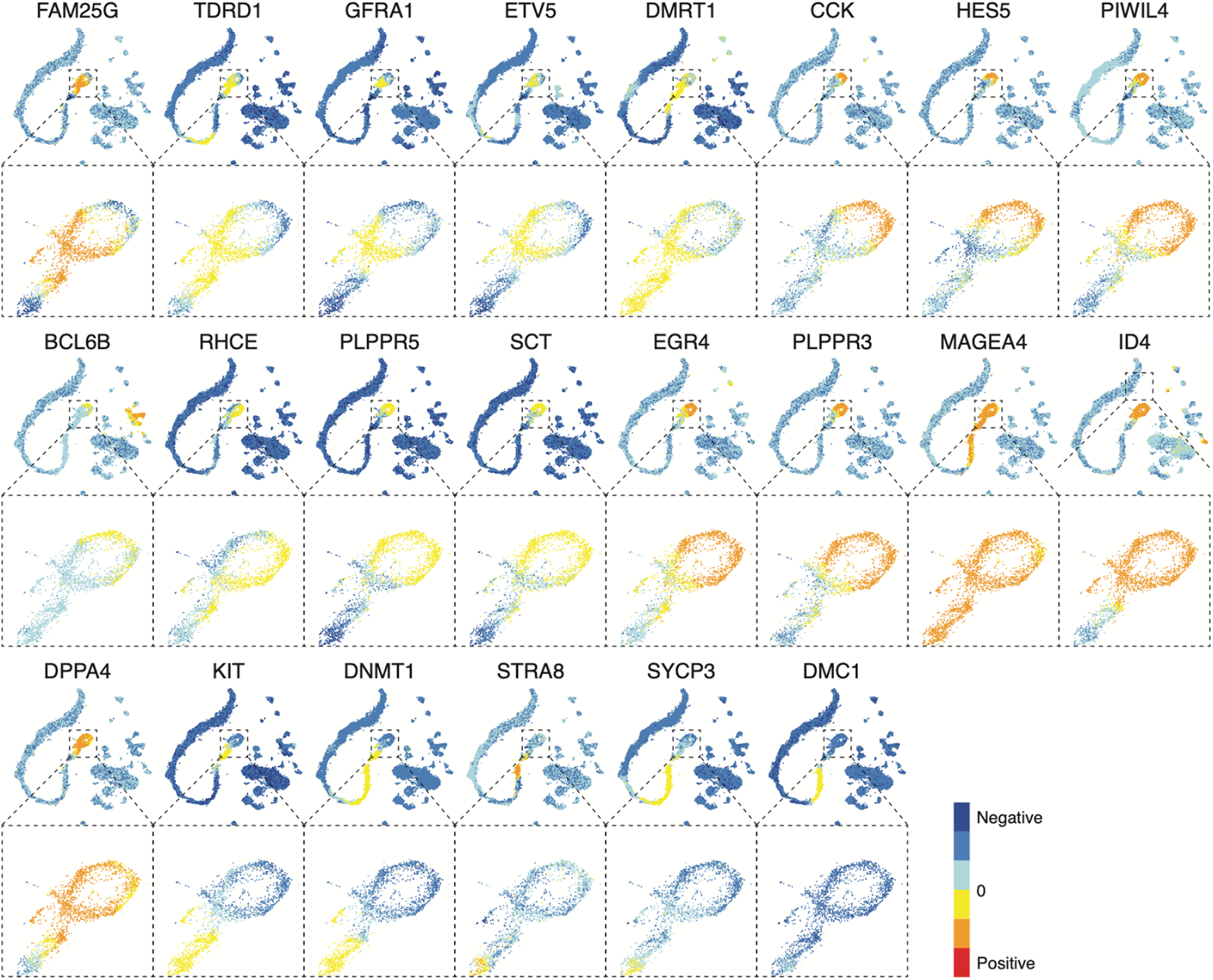
Expression levels found in HISTA of transcriptional markers of spermatogonial populations. These genes are found expressed in various patterns relative to the ring structure described in the main text; some span the entire ring (ID4), some also capture the stem (MAGEA4), whereas others have a precise localized expression (CCK, HES5, FAM25G, PIWIL4). Such patterns support the hypothesis that undifferentiated spermatogonial (USgs) cells in a cyclical continuum maintain the USgs population and feed spermatogenesis with new germ cells. Potentially, genes localized at the stem that also captures the attaching region of the ring (FAM25G, TDRD1, GFRA1, ETC5, and DMRT1) are candidates for future validation studies as checkpoint control of USgs terminating their maintenance cycle and entering spermatogenesis.

The USgs-associated components have the *NANOS* transcripts among several that cycle along the ring (Fig 6-7). *NANOS1, NANOS2,* and *NANOS3* collectively have not been previously described, as described herein, on human testis germ cells. *NANOS1* is not expressed as highly as *NANOS2* or *NANOS3* (Fig 6B), which is why it is not found to be highly loaded in any component but expressed in the *NANOS2* negative slice of USgs. Interestingly, on the 2D projection of the torus, *NANOS3* is expressed at the neck of the USgs ring structure, where the cell’s fate is decided; to cycle back as USgs or proceed with spermatogenesis. For a specific cell type quantification, HISTA’s ‘gene expression per cell type (boxplot)’ tab shows *NANOS3* is most explicitly expressed in early differentiating spermatogonial cells followed by other subsets of the spermatogonia. Furthermore, using the ‘Pseudotime gene” tab of HISTA, we can compare across pseudotime the exact populations which express these genes. Given these observations, we hypothesize that this structure is a biological signal suggesting a cyclical nature to these cells to maintain the stem populations and feed spermatogenesis with new germ cells.

## DISCUSSION

Single-cell transcriptomic analysis has become widespread in biomedical research, offering a powerful approach to unraveling cellular heterogeneity and identifying rare or elusive cellular populations and transcriptional signatures. Despite its immense potential, translating molecular findings in the high-dimensional space of single-cell data has posed significant challenges. To address such limitations, several tools and frameworks ^13,52,57^. These developments have paved the way for deeper insights and broader applications in single-cell transcriptomics. However, these software can mask the complexity of the data and computational methods and provide confounded results, as originally they were designed and parameterized with defaults for specific tissues or cell types such as PBMCs.

HISTA is a computational tool built on recent computational developments with the mantra to make a reference human testis and infertility scRNA-Seq atlas accessible for research while improving translational limitations to scRNA-seq. HISTA is unique compared to other websites or portals with testis scRNA-seq data; from its analytical approach to its data and represented cellular populations. HISTA enables a research environment that we plan to expand on and utilize as a basis to develop new analyses and future tools. Future planned developments include adding data from new donors, including cases of known forms of genetic infertility and samples to explore the developmental axis using fetal gonadal cells.

The results of our first computational study with HISTA provided novel insights into the regulatory processes of spermatogenesis, focusing on the role of lncRNAs and their potential implications for fertility. Our analysis revealed that a significant proportion of annotated lncRNAs show evidence of expression in the human testis transcriptome, suggesting their involvement in spermatogenesis. Identifying distinct lncRNA signatures in different transcriptional stages of spermatogenesis, from USgs, through meiosis, and to round spermatids, indicates their contributions to the maturation and function of the germ cells. Furthermore, as reported in the supplement of this manuscript, we found tight co-regulation between antisense and sense transcripts, which by pseudotime analysis elucidates new hypotheses about the transcriptional regulation of spermatogenesis. These findings are highly valuable for research and clinical perspectives, with possibilities for diagnostic and therapeutic application; this could involve targeting and manipulating lncRNAs, antisense transcripts, or their targets.

Next, we utilized the HISTA tool to investigate the hypothesis of USgs. We examined whether the ring structure observed in processing USgs data was a cyclical biological signal suggesting a fundamental mechanism for maintaining the stem cell population while continuously supplying new germ cells for spermatogenesis. The discovery of specific gene expression patterns and gene groupings, such as the NANOS family within the human USgs, provides insights into the regulatory mechanisms involved in SSC maintenance. Understanding the factors and processes that regulate SSC self-renewal and spermatogenesis can help develop strategies to enhance male fertility, such as contributing to advancements in assisted reproductive technologies and the preservation of fertility. As reported in supplemental work, we used the HISTA data with an alternate computational set of algorithms, validating and adding to our main findings with regards to USgs. However, further research is needed to elucidate the precise molecular mechanisms by which the identified genes, including the NANOS family, regulate SSC self-renewal and fate determination.

Nonetheless, there are still significant gaps in our knowledge of germ cell transcriptional regulation, including lncRNA-mediated regulation in human spermatogenesis. Firstly, precise molecular mechanisms of action remain largely unexplored. Additionally, the specific roles of such genes, especially lncRNAs in infertility and reproductive disorders, need further investigation, including their potential as therapeutic targets or predictive markers. Lastly, an important missing piece of the single-cell picture of the testis is the spatial landscape, connecting cellular heterogeneity with proximity, which we aim to be part of future releases of HISTA.

## DATA + CODE AVAILABILITY

HISTA code is available on GitHub (https://github.com/eisascience/HISTA), the processed R Shiny object can be found (https://conradlab.shinyapps.io/HISTA/), and the raw data used to generate this data can be found via our original manuscript.

## SUPPLEMENTARY CONTENT

We extend our results with two supplemental results sections. First as introduced earlier, we examine the transcriptional regulation of spermatogenesis by antisense transcripts. Next, we perform a computational validation investigating the cyclical structure of USgs with parallel methods that are mathematically distinct from our original approach housed by HISTA. Therefore, a supplemental section describes the application of SDA and its importance in HISTA. Lastly, we also provide a user manual for HISTA to guide users of specific capabilities; the most recent version of this manual can be found online as a tab within HISTA.

## AUTHOR CONTRIBUTIONS

JG, DTC, JMH, and KIA provided the surgical samples and most of the raw sequencing data. DFC and EM developed the analytical methods as well as HISTA. HISTA GUI and code was developed by EM. CD did an additional analysis on the USgs cells and contributed to completing the manuscript. ACL provided scientific consultation and performed validation experiments. KAV-C provided scientific consultation and developed illustrations and final figure panels.

## ACKNOWLEDGEMENT

With great appreciation to the National Institutes of Health (NIH), the LRP award committee, specifically the Contraception and Infertility Research from the Eunice Kennedy Shriver National Institute of Child Health and Human Development (NICHD) committee for supporting EM.

## SUPPLEMENTAL MATERIALS

### Supplemental Results: Transcriptional regulation by Antisense and sense genes

Antisense transcripts of the testis are another category of genes that can be explored as a set in HISTA through the ‘top loaded components’ tab. Similar to lncRNAs, we find several components enriched with antisense genes. Functionally, several proposed mechanisms exist by which antisense lncRNAs are considered to regulate sense mRNA ^58,59^. One mechanism involves forming RNA-DNA hybrids that are thought to stall the actions of RNA polymerase ^60^. Another suggested mechanism is the production of RNA-RNA hybrids, i.e., dsRNA, which can lead to siRNAs’ formation and the sense of mRNA’s degradation ^61^. More recently, with the discovery of the three-dimensional organization of the genome, certain lncRNAs have also been found regulating and organizing this structure, as well as nuclear organization and gene expression ^62^.

When investigating lncRNAs in our first vignette, we also observed some of the components to be enriched for antisense transcripts. Therefore using pseudotime analysis, we investigated the expression of these antisense transcripts relative to their sense-paired transcripts across spermatogenesis; a good example would be SDA109.

The expression pattern of the sense and antisense transcripts supports the hypothesis that these antisense transcripts play a role in regulating spermatogenesis. Using HISTA’s ‘pseudotime meta’ and ‘pseudotime gene’ tabs, the expression of such antisense transcripts and their matching sense transcripts can be visualized. For a select set of antisense and coding genes found in SDA109 across spermatogenesis pseudotime, we plot the expression of the antisense and sense pairs and the subtractive signal to illustrate a potential mechanism of transcriptional regulation (Sup Fig 1). Overall, we find that often where an antisense transcript is expressed, the coding transcript is low or undetectable. Furthermore, we observe that these antisense transcripts are frequently in adjacent transcriptional stages of spermatogenesis relative to the cells expressing the sense transcripts. As a specific and interesting example, Discs-Large Associated Protein 2 (DLGAP2) and DLGAP2-AS1 are two transcripts that have implications for the heritable disease.

*DLGAP2* is a membrane-associated protein expressed in the brain and the testis from SDA109, a case example illustrating the antisense-sene regulatory relationship. Only the paternal allele of *DLGAP2* is expressed in the testis ^63,64^. In HISTA, *DLGAP2-AS1* expression is localized at the transition between meiotically dividing spermatocytes and round spermatids. Such kinetics relative to the expression of *DLGAP2* suggests *DLGAP2* functions are inhibited in later stages of meiosis, and one of its regulators is likely *DLGAP2-AS1*. Alterations in *DLGAP2* have been linked to autism spectrum disorder and schizophrenia ^65,66^. Recent studies have identified this gene among several top candidates as having potentially heritable differentially methylated CpG sites in the sperm of cannabis-exposed men ^67^. Using similar deductive reasoning, other antisense and sense mRNA pairs related to spermatogenesis found in the same component (SDA109), such as *PPP4R1, JARID2, LNX1, TBX18,* and *SLIT1*, together with their antisense-mRNAs suggest a tightly regulated network. Such antisense expression is often overlooked in transcriptional analysis and was only found by using HISTA, because SDA has searched the entire detectable transcriptome for non-linear correlating manifolds in high-dimensional transcriptional space. Combined, the genes enriched in SDA109 and SDA52 highlight only a small section of a complex regulatory circuit of spermatogenesis, and the DLGAP2 highlights the potential for heritability of such tightly regulated and critical genes and their antisense counterparts.

### Supplemental Results: Validating Observations in Undifferentiated Spermatogonia with a parallel method

As described in the methods section, as a computational validation of our primary results obtained by using HISTA, we performed K-means clustering to identify an unbiased structure in gene expression related to USgs. We found the clusters identified by K-means do not divide USgs in a pattern that divides the donut structure; the clusters appear randomly distributed across the donut. Using a machine-learning classifier, we identified the most critical genes that explain this clustering, ranked by the Gini index (Supp Fig 2A). These genes are shown to be significantly differentially expressed across these clusters (Supp Fig 2B). Most of these genes overlap with those found by SDA. Interestingly, *NANOS2* and *NANOS3* are observed again as essential genes driving structure in the transcriptomic landscape of USgs.

Pseudotime analysis of USgs, given the K-means identified clusters, identifies a cyclical trajectory (Sup Fig C), which was used to compute differentially expressed genes across it, as depicted by a heatmap (Sup Fig D). Defining this trajectory’s actual start and end points is challenging, but we can predict the populations using several known markers discussed in the USgs vignette. For example, *STRA8*, found in differentiating spermatogonia, is not expressed in the donut and comes up later in the neck section; this can also be viewed in the ‘Pseudotime gene’ tab. *HES4*, a transcription factor found in several tissues, including the testis, is one of the earliest markers of USgs, and is highest in cluster 1 of the K-means clusters, whereas *NANOS3* is elevated in cluster 2. This suggests the ordering to be cluster 1, 3, 2, 4, which is the order that the heatmap pseudotime is ordered; starting halfway in the pseudotime, cells of 1 initiate this ordering pattern.

The fact that K-means clusters do not divide the donut like SDA components do, is inherent to the mathematics of each algorithm. K-means clustering minimizes the within-cluster sum-of-squares, i.e., seeking to find compact and well-separated clusters based on the distance between data points. In our case, the donut structure of USgs poses a challenge for K-means as these cells are not discrete populations, but rather a dynamic spectrum. Consequently, the algorithm needs help to capture the intricate 2D arrangement of USgs within the donut. On the other hand, SDA utilizes a probabilistic framework to identify coherent gene expression patterns and cells expressing them. This allows SDA to capture the circular structure exhibited by the USgs effectively. However, despite the divergent approaches, their results uncover underlying molecular factors shared between the two clustering strategies, providing converging evidence for the involvement of specific transcriptomic signals.

### SDA and scRNA-Seq data

SDA (Sparse Decomposition of Arrays) is a machine-learning algorithm that finds meaningful relationships in sparse, high-dimensional data; it is one of the unique core tenants of HISTA. SDA is a form of soft-clustering that has been demonstrated to be instrumental in untangling the complex scRNA-Seq landscape ^13^. Several key features of the SDA algorithm make it unique from other factorization methods^13^, mainly when applied to scRNA-Seq data^1,2^. In short, SDA helps to break down the 20,000+ detectable gene space into a manageable set of components, thus reducing the dimensionality of the original cells by gene matrix (DGE). This is in contrast to other pipelines that initially reduce the gene space to 2000-3000 top detectable genes prior to a linear (PCA) dimensionality reduction. In our application, SDA inputs the DGE and outputs two sparse matrices: the gene loadings matrix (genes by component) and the cell scores matrix (cells by components). By definition, their dot product reproduces the original gene expression matrix. The key takeaway is that SDA finds relationships that go beyond the current pipelines of scRNA-Seq (found in almost all other web portals of scRNA-seq data) that rely on performing differential expression (DE) analysis on clusters identified in a space defined by a heuristically reduced set of principal component. Another key takeaway from component annotations is that each SDA component scores the cells in HISTA. This score represents a spectrum in which the cells relate to the function of that component.

### HISTA User Manual

#### Tabs

##### Home Page

The Home page of HISTA provides a graphical introduction to navigating HISTA. On the left is a tabbed menu bar to navigate the main features of HISTA, described in detail below.

##### Main tab

This is the ‘Main’ tab that HISTA loads. On the center top, two information boxes provide available background information for the selected SDA component and gene found in the interactive menu on the left panel of this tab. This menu has several parameters to change that alter what is being displayed.

From top to bottom of the page:

● In the Inputs sections, the first input allows searching SDA components via a numerical input.
● In the next box, genes (human symbols e.g. PRM1) can be searched to display the gene’s t-SNE projected expression. Additionally, HISTA will highlight the components that the gene of interest is mostly found or, in other words, highly loaded.
● In the next selection, via the radio button provided, it is possible to select which of the pre-processed t-SNE plots are shown (there are four options):

● The batch-removed SDA cell score matrix t-SNE (default). This t-SNE was produced on non-batch components of the SDA score matrix i.e., cells by components
● The batch-removed SDA cell score matrix UMAP.
● The pre-SDA DropSim-normalized expression matrix t-SNE. This t-SNE was produced on the gene expression matrix normalized by DropSim, which includes a square-root-based transformation.
● the batch-removed SDA-imputed expression matrix t-SNE. As explained earlier, by performing dot-product of the gene loading and cell score matrices, with only the non-batch components, we impute a batch-removed gene expression matrix on which t-SNE was run.
● The next set of radio buttons is to visualize available metadata such as donors, replicates, conditions, experiments, cell cycle, and cell type (default).
● The final items in the interactive menu are several buttons to download the top-loaded gene lists for export as well as manual navigation of the SDA components.
● The figures below the input section from top to bottom include

● the gene expression, cell score, and metadata 2D projection (t-SNE or UMAP).
● The cell score across donors scatter plot, which aids in seeing the selected component’s score distribution
● The gene ontology (GO) plots proved a pre-computed analysis of the top loaded genes that aid in translating the general signature observed.
● The chromosome location highlights the loading weight of each gene relative to their position across the chromosomes.
● At the bottom of this tab, the top loaded positive or negative genes are ranked and listed, but how many are shown can be inputted, with 20 being the default.

##### Index of Components

A table of the SDA components and our observations summarized pertaining to each. To curate this table, iterative rounds of analysis were performed on each component, representing a summarized form of our SDA findings.

##### Fingerprinting (heatmap)

Two heatmaps, identifying signature/barcode pattern, annotating each component quantitatively, one for positive and one for negative cell scores. Chi-squared analysis of the number of cells (positive or negative) per component identifies the enrichment of these cells per component. When contrasted by pathology (CNT, INF1, INF2, KS, JUV), we add an extra dimension to the enrichment analysis. The pair-wise hierarchical clustering (which can be turned off using the radio buttons provided) identifies similar enrichment patterns for positively or negatively scored cells, split by available metadata options, selectable via radio buttons, e.g., cell types, donors, pathology, etc. The columns represent the SDA components, and the rows are the experimental conditions. By defining thresholds on the cell score matrix, the number of cells that are scored positively or negatively are enumerated and passed through a Chi-Squared test; visualized are the transformed residuals that highlight enrichment or depletion by performing pairwise hierarchical clustering, the components and the samples that are most similar, cluster together. Interestingly, one of the results of this analysis supports our other findings that INF2, the patient with secondary azoospermia, is more similar to the adult controls than INF1, the idiopathic azoospermia patient.

##### Gene Expression per pathology (boxplot)

Gene expression boxplots of a searched gene across pathology (CNT, INF1, INF2, KS, JUV) with Wilcox testing, with the ability to subset by cell type. Specifically, to test if the expression of a single gene is distributed similarly (computing p-value via Wilcox-rank-sum test) between the available experimental conditions in HISTA (CNT, KS, JUV, etc.); additionally, through the available radio buttons, it is possible to focus the hypothesis testing by cell type. For example, we can search XIST across all cell types and find it is significantly enriched in Klinefelter Syndrome (KS) (p<2.22e-16 compared to controls). Next, by selecting Sertoli cells (SC), we find that this significant enrichment is lost, as previously reported^1^.

##### Gene Expression per cell type (boxplot)

Gene expression boxplots of a searched gene, across cell types, with the ability to subset by pathology (All, CNT, INF1, INF2, KS, JUV). This tab enables the user to quantify the expression of a specific gene across all cell types. Unlike the per pathology boxplots, statistics are not provided due to minimizing the complexity of the figure. For example, we can observe that the Sertoli cell marker SOX9 is exclusively expressed in Sertoli cells. This approach is powerful at rapidly visualizing expression distribution across the available cell types.

##### Gene Expression per cell type (2D)

Gene expression of a searched gene, batch-corrected, mapped on the t-SNE projection, with the ability to subset by cell type. Additional radio buttons allow selecting which 2D plot to show, e.g., t-SNE or UMAP, on various scopes of our data.

##### Cell scores per cell type (2D)

Scores of a searched component, mapped on the 2D (t-SNE or UMAP) projection, with the ability to subset by cell type. To provide a deeper scope of visualizing the cell scores projected on the pre-computed 2D representation. In this tab, it is possible to subset the figure by cell type using the available radio buttons. For example, SDA component 1 demonstrates that the highest absolute scores are found in the spermatid population of the germ cells. By zooming in on these cells, we can better observe the distinct banding pattern that positively scores the last and early spermatids but negatively scores the spermatids between them. Digging into the gene loadings of this component, we observe specific gene regulation patterns that explain the banding observed in the cell scores; for example, the top positively loaded genes contain *SPRR4* and *PRM1*, whereas the top negatively loaded genes contain *FSCN3* and *PRM3* supporting the regulation of spermiogenesis as the spermatids finish their maturation.

##### Metadata per cell type (2D)

In this tab, available metadata (selectable) are mapped on a 2D projection which can be subset by cell type (selectable). This tab enables the user to create parallel visualization to the focused t-SNE or UMAP of the cell score or gene expression 2D tabs, but with metadata, providing a closer look at the origin and background of each cell.

##### Gene correlations

In this tab, the user can input a set of genes and select a cell type via the radio buttons, which visualize a heatmap of gene-gene correlations within the selected cells.

##### Component Correlations

The top-loaded genes really are at the core of many assessments, including the translation of findings. We provide a way to explore and evaluate the relationship of the components narrowed by the number of top genes. First, select a component of interest (numeric input). Then use the slider to select how many top genes to include in the correlation analysis. Visualized is a heatmap of component-component correlation. For convenience, the top-loaded genes used are shown.

##### Pseudotime Meta (germ only)

A pseudotime trajectory, as previously described, was inferred on the t-SNE 2D projection of the germ cells, giving order to cells driven by the transcriptomic landscape that also parallels the known spermatogenesis trajectory. Plotting the cell scores of SDA components (y-axis) pertaining to germ cells across the cells ordered by pseudotime (x-axis), we commonly see “wave” patterns that translate to transcriptional kinetics of that component relative to the affected cells. There is an additional ability to select available metadata to identify correlating scoring patterns.

##### Pseudotime Gene (germ only)

This tab allows users to type in a gene of interest to visualize the expression wave across our defined pseudotime. Additionally, metadata selection is available to visualize differential expression patterns.

##### Pseudotime Component Index

An index of the SDA components pertaining to germ cells and a summary of our observations. A key feature of this table is the order given to these components relative to the cells (i.e., stages of spermatogenesis) they score with the most magnitude. This ranking is found by identifying the main peak/maxima of the density curve that is fitted to the scores (y-axis) by pseudotime (x-axis) scatter via a peak finding algorithm; the position of this maxima defines the rank. For example, the third-ranked SDA component, SDA #149, splits early spermatogonial cells into 3 sections where the intermediate section is scored negatively opposite to the others, which we describe more in detail in the vignettes section.

##### Enrichment analysis

Given a set of genes, hypergeometric testing (K=150) identifies which components are highly enriched with those genes. In application, we find an adjusted p-value less than 0.01 to find significant enrichments. However, the fold enrichment can also be highly informative when significance is not determined. This statistical approach is most appropriate for less than ∼30 more than 3 genes.

##### Top loaded components

This tab is another statistical approach to identify key components that parallel enrichment analysis, although this method is best suited for larger sets of genes. As a default, the input is loaded with antisense genes. There are several figures in this tab. First, there is a histogram layered with a density plot comparing two distributions. Given a set of genes enriched in some biology of interest, e.g., antisense genes, we can compute the distribution of how many of these genes are found across the top positive and negative loaded components. This distribution is layered over a second distribution derived from an equal-length random sample of genes (computed dynamically). By comparing these distributions, we can evaluate components enriched for or depleted of our selected genes relative to a random sample.

In the next plot, the components are ranked by how many genes of the input set are found in the top loaded components, split by positive and negative loading direction. Based on the first figure in this tab, we can use general statistics like the mean number of genes derived from a random set (∼9 for the default gene set) to filter which components are more or less than average enriched or depleted respectively of the input gene set.

The last figure is a correlation heatmap of the gene loading components filtered by the input gene. This identifies structural similarity amongst the components in the defined gene space. Each component is enriched with a certain pathology and cell type annotated on this heatmap’s rows.

##### lncRNAs

As described in this manuscript’s long non-coding RNA (lncRNA) vignette, we have reproduced some of these figures. Additionally, this tab yields the lncRNAs are provided by entering the component number interest in the input box. On the top, a Venn diagram shows the overlap of Ensembl lncRNA annotated genes and all genes found in HISTA. As described in the ‘top loaded components’ tab, we observe distribution overlap of the number of these genes found in the top loaded genes across components, compared to a random equal-length set of genes. Then the components are sorted and split by direction to identify lncRNA-enriched components.

##### Soma only W. LN19

In 2019 a new Klinefelter scRNA-Seq was made available through collaboration with Leurentino et al., but after our initial data freeze, the SDA-HISTA analysis. Furthermore, after integrating this new single KS donor using our customized Seurat pipelines, we found evidence for differences that we believe are correlated with the sampling procedure and the age and health of the donor. Therefore, we plan to combine this data in later releases of HISTA. However, we limited our analysis to only validating our results with this new data for the current scope and release. This tab is provided as support material for our KS manuscript1.

##### Brief notes

● As with the existing KS donors, there were no germ cells in this new patient, so we could only focus on the somatic cells.
● There is a single large cluster of Sertoli cells (SC) comprising at least 3 sub-clusters.

○ The two smaller in diameter subclusters are enriched with JUV SC, but only one is enriched with the SC from other adults.
○ The largest SC subcluster is mostly derived from LN19.
We previously defined 3 major subtypes of Leydig cells (LC); the progenitors (PLC), the immature (ILC), and the mature (MLC) groups.

○ The LN19 LC are found mostly in PLC and MLC clusters.
● LN19 contributes a distinct cluster of cells that we believed to be of myoid phenotype.

##### LC only W. Zhao21

In 2021, a new scRNA-Seq dataset was published by Zhao et al. We plan to combine this data in later releases of HISTA; however, for the current scope and release, we limited our analysis to only validating our results with this new data. More recently, in attempts to download the raw sequencing files of this data, significant anomalies have been identified, which may limit our future integration of all of this data. Many of the samples from this dataset failed bioinformatics QC processes. Nonetheless, using their processed count data, we validated our Leydig cell findings as support material for our KS manuscript^1^. As mentioned above, we have previously defined 3 major subtypes of Leydig cells (LC); the progenitors (PLC), the immature (ILC), and the mature (MLC) groups. By combining the LN19 and Zhao21 data, we validate we find these clusters in an integrated dataset composed of cells from multiple donors, which is critical as these populations, such as the MLCs, are fairly infrequent relative to other cells of the testis, especially in normal conditions.

##### HISTA Case Examples

<u>Case example 1:</u> You have one or more genes, and wish to learn more about them within HISTA.

● To look at the expression of each gene, search for the gene in the main tab

● The figure header of the 2D (t-SNE or UMAP) expression plot in the main tab also lists in order of the SDA components and the associated loading, which enables identifying gene sets that correlate with the gene of interest
● To test pathology differential expression, on specific cell type (or all cells), the ‘gene expression per pathology’ tab visualizes this with Wilcox statistics.
● To visualize expression across cell types but in specific pathologies, use the ‘gene expression per cell type’ tab.
● If this gene is a germ cell expressed gene, try the ‘pseudotime expr’ tab to observe the expression pattern across pseudotime.
● If you have more than ∼3 genes, you can use the ‘Enrichment analysis’ tab to find which components they are enriched in and search those components for annotation and find additional correlating genes. You can also use the ‘top loaded components’ tab if your set of genes is fairly large.

<u>Case example 2:</u> You have a cell type of interest, and you wish to learn about the genetic signatures found within HISTA pertaining to the cell type.

● Identify the location of the cell type of interest in the ‘main’ tab.
● Using the component index, identify which components score to this cell type and search them in the ‘Main’ tab to identify the expressed genes.
● Use the ‘Gene expression per cell type’ tab to evaluate cell type specific expression

<u>Case example 3</u>: You have a hypothesis about a gene/transcript, and you wish to see if it is differentially expressed across experimental conditions found in HISTA.

- ○ Use the ‘Gene expression by pathology boxplot’ tab to identify significant differences

#### Supplemental Figures

**Fig Supplement 1:**
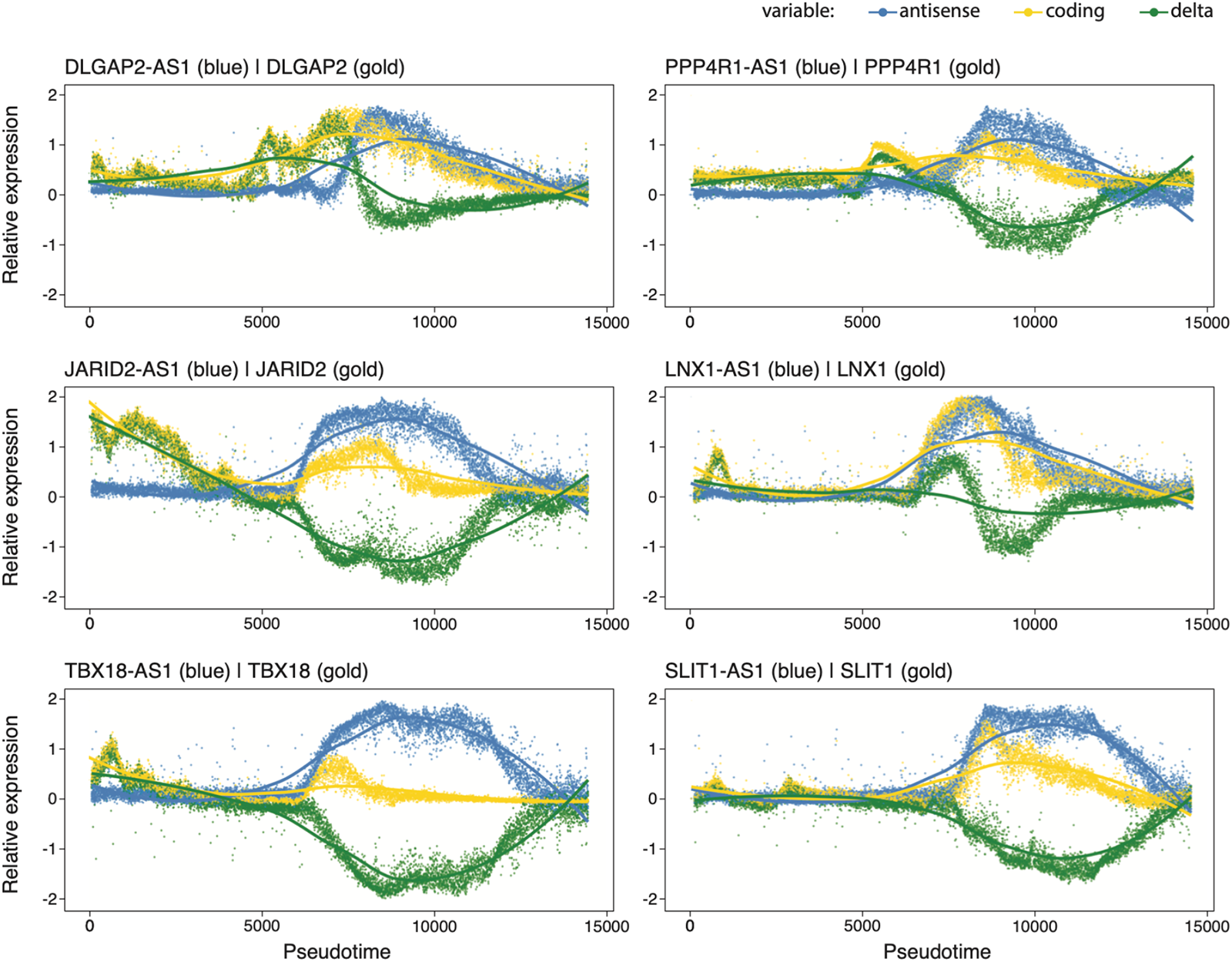
Antisense and sense transcripts demonstrate tight regulation in transcriptional regulation across spermatogenesis. Shown on the y-axis are the expression levels of select antisense genes (blue) and their matching sense pairs (gold), as well as their delta defined simply as sense minus antisense (green). This set is specifically derived from component 109 of HISTA, which targets spermiogenesis. Antisense expression is often not part of the discussion in sc-RNAseq analysis; however, in such critical contexts such as transcriptional regulation of germ cells, the delta expression highlights an alternative kinetic regulation by these genes and their antisense partners.

**Fig Supplement 2:**
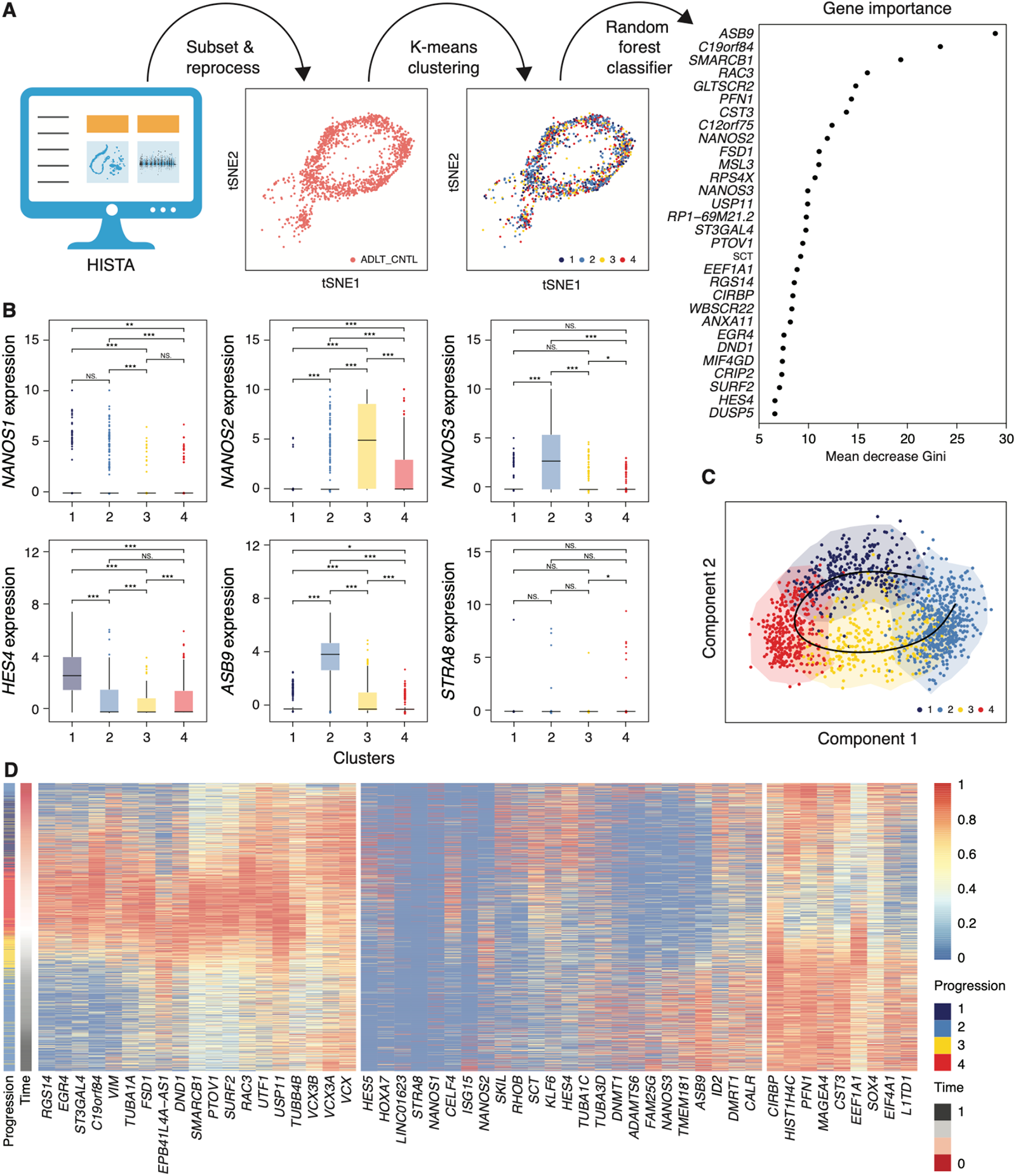
We performed parallel computational validation to our main analytical framework of quantifying the transcriptional landscape of undifferentiated spermatogonia (USgs). **A)** Using the HISTA dataset, we subset only for USgs and reprocess the data (see Methods). Briefly, we performed unsupervised K-means clustering on the gene expression of the top 500 variable genes in USgs, which unraveled at least 4 clusters. Using a Random-forest classifier, training on the K-means derived clusters, we identify the top most important genes that explain this structure as depicted by a Gini Index. **B)** NANOS and select genes of high importance are shown to be significantly expressed by the K-means clusters (Wilcox, NS = not significant, * adj.p < 0.01, ** <0.001, *** < 0.0001, and **** < 0.00001). **C)** Pseudotime analysis identified a trajectory across the K-means clusters using the same 500-top-variable gene space, which is found to be cyclical as depicted by the gene expression patterns in **D)** as the pseudotime-ordered gene expression heatmap. Note the progression pattern starts and ends with cluster 2 (light blue) that parallels the cyclical pseudotime derived in **C**.

